# Real-time HRMAS ^13^C NMR of obligately anaerobic cells identifies new metabolic targets in the pathogen *Clostridioides difficile*

**DOI:** 10.1101/2021.05.04.442336

**Authors:** Aidan Pavao, Brintha Girinathan, Johann Peltier, Pamela Altamirano Silva, Bruno Dupuy, Leo L. Cheng, Lynn Bry

**Author notes:** Corresponding authors: Lynn Bry, MD, PhD;, Leo L. Cheng, PhD.

## Abstract

Anaerobic microbial metabolism drives critical aspects of host-microbiome interactions and supports many economically important industrial applications. Yet, the metabolic pathways of anaerobic bacteria and their associated constraints for maintaining energy and redox balance are often poorly described. We employ High-Resolution Magic Angle Spinning Carbon-13 (^13^C) Nuclear Magnetic Resonance spectroscopy with dynamic flux balance analysis to resolve real-time metabolism in living cells of the obligately anaerobic pathogen *Clostridioides difficile*. Using ^13^C-labeled carbon sources, we elaborate the time-dependent progression of reductive and oxidative anaerobic fermentation pathways. Analyses identified new integration points for redox and nitrogen coupling between carbohydrate and amino acid metabolism, particularly in the production of ^13^C-alanine from ^13^C-glucose to provide an ammonia sink from co-occurring amino acid fermentation. Analyses conducted in the presence or absence of selenium, a required co-factor for the proline Stickland reductase, demonstrate further capacity to modulate cellular metabolism and resulting metabolites. Findings informed a genome-scale metabolic model of *C. difficile*, identifying alanine and associated electron carrier pools as critical metabolic integration points in energy flow and biomass expansion. We illustrate use of HRMAS NMR as a new analytical platform to resolve complex interactions in anaerobic metabolism and inform new metabolic targets to counter *C. difficile* infections.

## Introduction

Obligate anaerobes influence crucial aspects of human health and disease. These species form the majority of members of the mammalian gut microbiota [1] and include pathogenic species such as *C. difficile*. Anaerobic bacteria also have many important industrial applications, including the production of solvents, biofuels [2, 3] and foodstuffs [4]. However, the study of obligate anaerobes poses unique challenges given the need to maintain anaerobic culture conditions. Moreover, the metabolic pathways and nutrient requirements of obligate anaerobes often differ substantively from those of model species such as *Escherichia coli* or *Bacillus subtilis*. Thus, despite their clinical and industrial importance, many anaerobes remain poorly characterized.

High-Resolution Magic Angle Spinning (HRMAS) NMR is a powerful tool for the metabolic study of complex samples [5] including of living cells [6]. HRMAS studies of continuous *in vivo* metabolism by NMR (CIVM-NMR) have successfully resolved real-time metabolism in human leukemia cells and in the ascomycete fungus *Neurospora crassa*, including in media supplemented with uniformly-labeled ^13^C compounds [7]. HRMAS NMR is particularly suited to the study of anaerobe metabolism as the rotor chamber can maintain an anaerobic environment [7]. However, use of HRMAS NMR with ^13^C-labeled substrates has not previously been applied to study time-dynamic metabolism in obligate anaerobes.

The obligately anaerobic pathogen *Clostridioides difficile* is classified as an urgent threat by the U.S. Centers for Disease Control and Prevention [8]. The pathogen causes over 450,000 infections and 20,000 deaths annually in the US and is the leading cause of hospital-acquired infections [8, 9]. *C. difficile* readily colonizes gut environments through its capacity to ferment diverse carbon sources including carbohydrates, complex polysaccharides, amino acids, and ethanolamine [10]. *C. difficile* possesses multiple amino acid fermentation enzymes including the Stickland glycine and proline reductases, both of which require selenium for activity [11–13], and the reductive leucine pathway which is not selenium-dependent [14]. The pathogen can also fix carbon dioxide through the Wood-Ljungdhal pathway to generate acetyl-CoA for further metabolic and biosynthetic reactions [15, 16].

A complex network of gene regulatory and metabolic programs integrates *C. difficile’s* virulence with its metabolism [17, 18]. As growth of *C. difficile* exceeds the carrying capacity of gut environments, the organism releases toxins as a means to obtain nutrients from damaged host tissues [8, 19]. A deeper understanding of *C. difficile’s* real-time metabolism opens opportunities for new approaches to prevent and treat infections, including ones that do not rely solely on antimicrobial therapy [18]. However, achieving this goal requires accurate information regarding the pathogen’s complex metabolism and its integration with cellular processes modulating energy balance, growth, and virulence.

Metabolic models for *C. difficile* provide an additional important tool to frame observations from NMR studies in the context of metabolism at the cellular scale. NMR methods incorporating ^13^C-labeled substrates have long been applied to constrain steady-state metabolic flux analysis [20–22]. Dynamic flux balance analysis (dFBA) approaches using input time-series from GC-MS metabolomic data have also been used [23] to model *C. difficile* metabolism [24].

Using HRMAS NMR with uniformly labeled ^13^C carbon substrates, we tracked the molecular context of ^13^C atoms to resolve *C. difficile’s* metabolic pathways. dFBA analyses, informed by HRMAS ^13^C NMR findings, elucidated active metabolic pathways and associated gene targets. Our approach validated known pathways of carbohydrate and amino acid fermentation, described the temporal recruitment of metabolic systems, and uncovered new integration points between glycolytic and amino acid metabolism, including critical processes maintaining cellular redox and nitrogen balance. We demonstrate integrated application of HRMAS ^13^C NMR with dynamic flux balance analysis to resolve obligate anaerobe metabolism and inform targeted functional applications.

## Results

### HRMAS NMR of ^13^C-glucose identifies time-dependent changes in *C. difficile* metabolism and new mechanisms for maintaining redox and nitrogen balance

We utilized HRMAS NMR’s technical advantages to obtain high resolution proton and ^13^C NMR spectra from living cells. *C. difficile* vegetative cells were added to Modified Minimal Media (MMM; [25]) prepared with required natural abundance carbon sources and a defined ^13^C carbon source (Methods, Table 2). The use of uniformly ^13^C (U-^13^C) labeled substrates supports definitive tracking of complex metabolic pathways by following the molecular context of ^13^C atoms.

Metabolism of U-^13^C glucose (Figure 1A-B, dark blue plots, Table 1, Supplemental Figure 1) over 48 hours identified rapid production of ^13^C-acetate from oxidative metabolism of ^13^C-glucose (Figure 1A-B, red plots), followed by ^13^C-carbonic acid by 3.9 hours (H_2_CO_3_; Figure 1A-B, light blue plot), representing dissolved CO_2_. By 4.9 hours of analysis, levels of ^13^C-alanine rose rapidly (Figure 1A-B, green plots). At 11.5 hours, ^13^C-CO_2_ was detected (Figure 1A-B, yellow plots). ^13^C-lactate and ^13^C-ethanol were detected at 13.4 hours (Figure 1A-B, black and pink plots, respectively) from reductive metabolism of ^13^C-pyruvate and ^13^C-acetate, respectively, with oxidation of NADH to NAD+ [16]. ^13^C-pyruvate signal remained undetectable, suggesting its rapid conversion to the observed metabolites.

**Figure 1:**
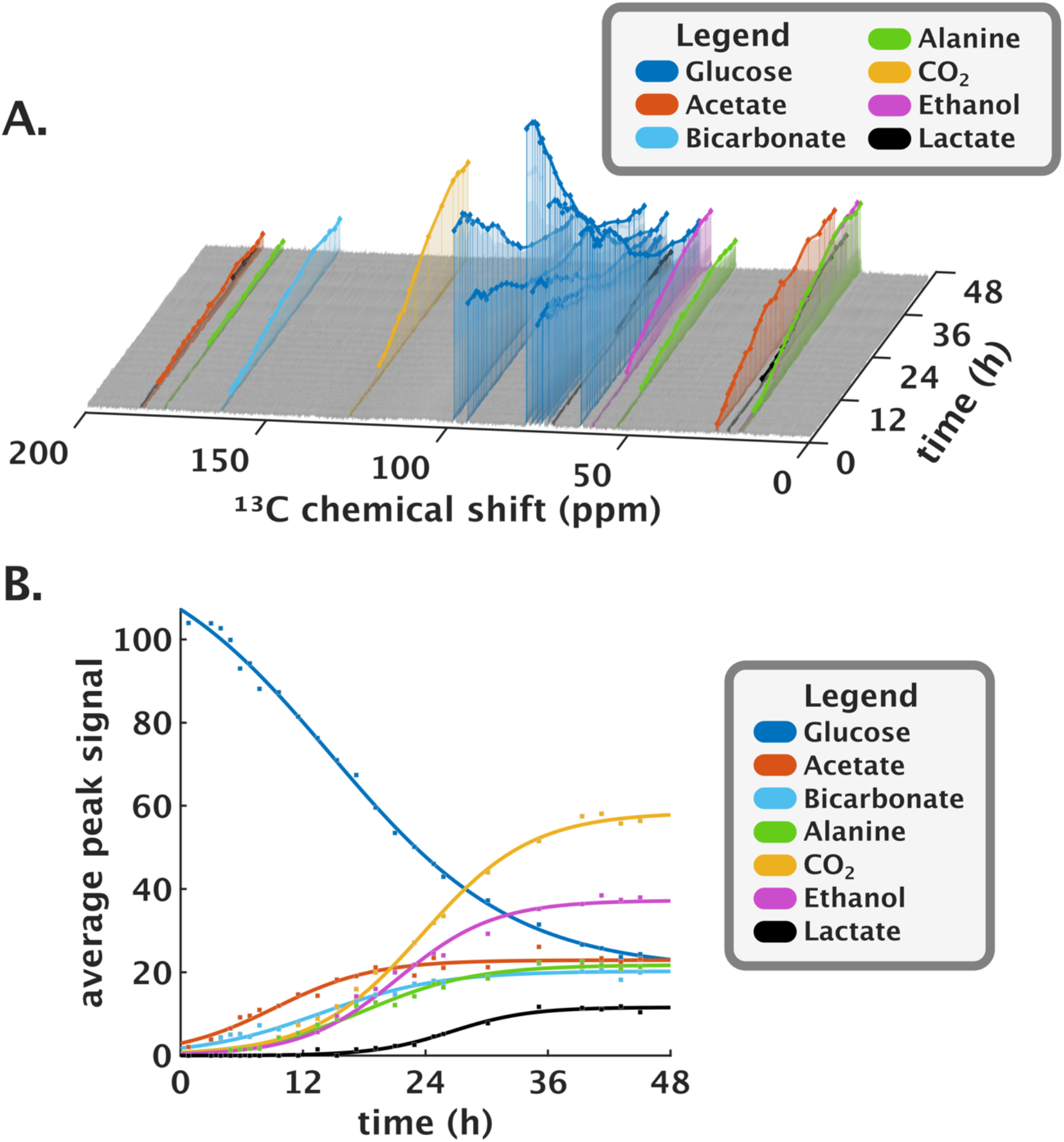
HRMAS ^13^C NMR resolves real-time metabolism of U-^13^C glucose by *C. difficile*. (A) ^13^C-NMR spectrum time series stack plot. X-axis shows the ^13^C chemical shift; Y-axis the time in hours, and Z-axis the ^13^C normalized signal. Inset key indicates the signal for individual carbons within the detected compounds. (B) Average peak signal of detected ^13^C compounds (Y-axis) over time (X-axis) and logistic curve fit. Individual points were normalized by the number of ^13^C carbons per compound.

**Table 1.** Kinetic parameters of ^13^C-labeled compounds detected by HRMAS ^13^C NMR.

|  |  | Logistic function coefficients |  |  |  |  |
| --- | --- | --- | --- | --- | --- | --- |
| Metabolite | Detection Time (h) | Upper bound $L$ | Rate constant $k$ (h<sup>-1</sup>) | Maximum growth rate $kL/4$ (h<sup>-1</sup>) | Half-maximal time $x_0$ (h) | Final signal $C$ |
| U-<sup>13</sup>C-Glucose +Selenium |  |  |  |  |  |  |
| Glucose | 0 | 103.6 | −0.111 | −2.87 | 14.7 | 20.5 |
| Acetate | 0 | 22.9 | 0.207 | 1.19 | 9.2 | — |
| Bicarbonate | 3.9 | 20.3 | 0.171 | 0.87 | 13.7 | — |
| Alanine | 4.9 | 21.8 | 0.185 | 1.01 | 18.1 | — |
| CO2 | 11.5 | 58.5 | 0.187 | 2.73 | 23.7 | — |
| Ethanol | 13.4 | 37.3 | 0.213 | 1.98 | 21.4 | — |
| Lactate | 13.4 | 11.6 | 0.264 | 0.77 | 26.6 | — |
| U-<sup>13</sup>C-Proline +Selenium |  |  |  |  |  |  |
| Proline | 0 | 78.8 | −0.261 | −6.86 | 8.9 | 46.6 |
| 5-aminovalerate | 2.5 | 59.0 | 0.419 | 8.24 | 9.8 | — |
| U-<sup>13</sup>C-Proline −Selenium |  |  |  |  |  |  |
| Proline | 0 | 70.0 | -0.187 | −3.267 | 2.95 | 69.9 |
| 5-aminovalerate | 12.0 | 8.6 | 0.183 | 0.39 | 21.03 | — |

### Reductive proline metabolism co-occurs with reductive glucose metabolism

*C. difficile’s* Stickland proline reductase drives energy production and growth through the paired metabolism of electron-donor amino acids such as branched-chain amino acids, alanine and aromatic amino acids, with the electron recipient proline [11, 26]. In contrast to the glycine reductase system, the proline reductase directly couples to the membrane-bound Rnf electron transport system and has been shown to efficiently drive ATP production and cellular growth [27].

In media conditions containing U-^13^C-proline, the Stickland proline metabolite 5-aminovalerate appeared within 2.5 hours and attained maximum signal by 20 hours, representing consumption of 60% of ^13^C-proline (Figure 2A). Production of ^13^C-5-aminovalerate occurs only via the proline reductase enzyme with no subsequent metabolism by *C. difficile*. The maximum rate of 5-aminovalerate production was 10.5% per hour of the maximum signal (Figure 2C, Table 1).

**Figure 2:**
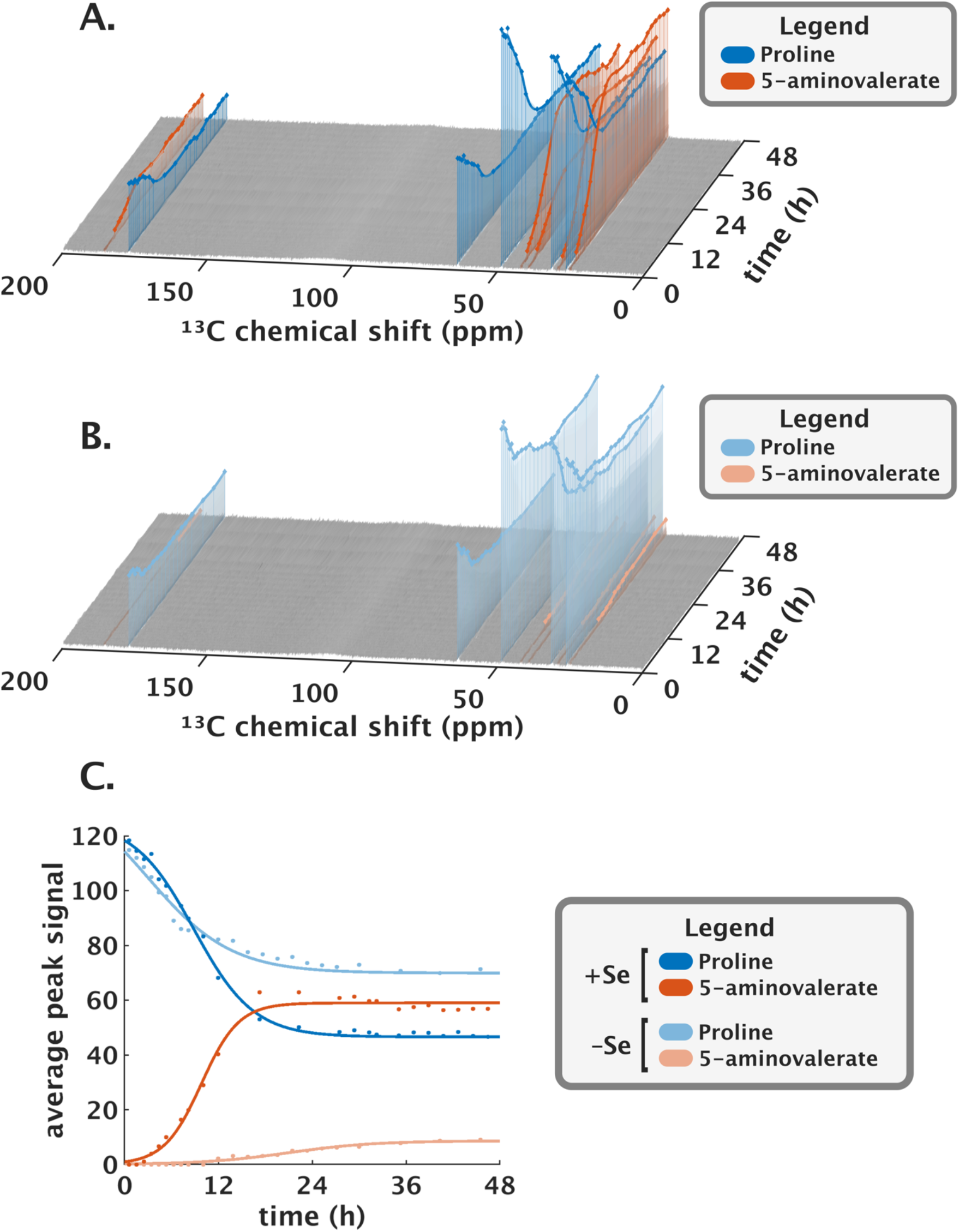
HRMAS ^13^C NMR reveals dependence of proline metabolism on selenium concentration. Surface plots of ^13^C-NMR spectra for reactions containing U-^13^C proline with (A) 0.1 μM sodium selenite or (B) in selenium-deficient media. (C) Average peak signals over time of proline and 5-aminovalerate in trace and 0.1 μM sodium selenite reveal increased proline utilization and 5-aminovalerate production in the presence of non-limiting selenium.

### Exogenous media conditions modulate Stickland proline metabolism

To further evaluate HRMAS NMR’s capacity to detect altered cellular metabolism by modifying the exogenous media conditions, we measured the real-time cellular metabolism of U-^13^C-proline under conditions of limiting selenium (Figure 2B). In the absence of exogenous selenium, ^13^C-5-aminovalerate’s detection was delayed to 12 hours of analysis and showed a 20-fold reduced rate of production, with maximum levels reduced by 85% as compared to media containing selenium (Figure 2C, Table 1).

### Improved *C. difficile* metabolic model informed by HRMAS NMR studies

We used the experimental quantification of the metabolites from ^13^C-glucose and ^13^C-proline metabolism to further enhance icdf834, a genome-scale metabolic model (GEM) of *C. difficile* (Figure 3) [28]. Model updates included addition of corresponding redox and energy generating reactions in glycolytic and Stickland metabolism (Supplemental Tables 1 and 2). The integrated signal of homologous carbons among molecules, fit to logistic functions, was used to predict relative ^13^C metabolite concentrations over time. Uptake and secretion bounds within the model were then set to the rates of change of the predicted experimental ^13^C metabolite concentrations.

**Figure 3.**
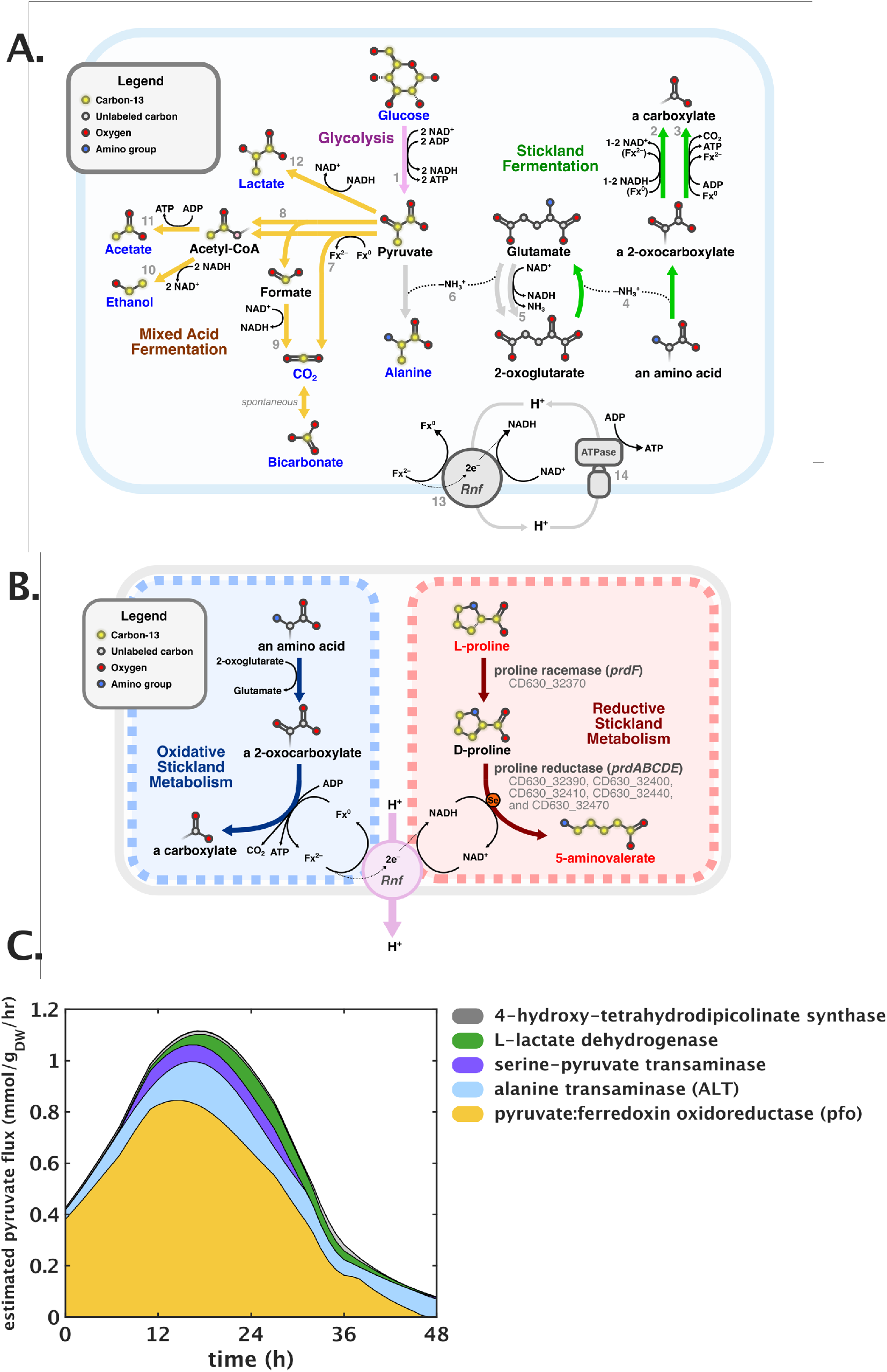
Metabolic pathways of *C. difficile* explain the time-dependent modulation of metabolism observed in HRMAS ^13^C NMR studies. (A) Glucose metabolism incorporates glycolysis and mixed-acid fermentation and is redox- and nitrogen-coupled with Stickland fermentation. Greyed numbers indicate specific reactions and associated genes in Supplemental Table 1. Labels of detected ^13^C-labeled metabolites are highlighted in blue. (B) Reductive proline Stickland metabolism is dependent on the *prd* operon genes, including the selenoenzyme proline reductase, and is redox-coupled to oxidative Stickland pathways via the Rnf complex. Labels of detected ^13^C-labeled metabolites are highlighted in red. (C) dFBA predictions of reactions driving pyruvate flux. X-axis shows time in hours, Y-axis shows predicted pyruvate flux in mmol/gram of dry weight biomass per hour. Inset key indicates the enzymatic reactions predicted to carry ≥1% of pyruvate flux.

Dynamic flux balance analysis (dFBA) solutions of the updated metabolic model predicted the expected conversion of glucose to pyruvate via glycolysis (Supplemental Tables 1, 2), with pyruvate:ferredoxin oxidoreductase (*pfo*) as the primary enzyme oxidizing pyruvate to acetyl-CoA and CO_2_. Flux through *pfo* accounted for ≥50% of total pyruvate flux over the first 40 hours of metabolism (Figure 3C). In contrast, pyruvate formate-lyase (*pflABCDE*), which metabolizes pyruvate to acetyl-CoA and formate, was not predicted to carry significant flux under these culture conditions. Downstream, the dFBA solutions indicated ≥75% of acetyl-CoA generated from pyruvate to be secreted as acetate. Acetyl-CoA’s oxidation to ethanol was predicted to peak at 24 hours, representing 10% of acetyl-CoA flux. The remainder of acetyl-CoA was predicted to be consumed in other biosynthetic pathways.

Transamination of pyruvate to L-alanine was predicted to constitute 9% of pyruvate flux initially, rising to 23% by 20 hours and 89% by 48 hours. The majority of L-alanine-forming transamination was predicted to occur via alanine transaminase (*alt*), using glutamate as the amino donor (Figure 3C). However, at 20 hours, the peak of L-alanine formation, 27% of alanine-forming transamination reactions were predicted to utilize serine as the amino group donor via serine-pyruvate transaminase (CD630_09940; Figure 3C).

Pyruvate flux through other pathways was predicted at lower rates. The dFBA solution estimated lactate production to peak at 30 hours, representing 12% of total pyruvate flux at this timepoint. Assimilation of pyruvate into 4-hydroxy-tetrahydrodipicolinate by 4-hydroxy-tetrahydrodipicolinate synthase (*dapA*, CD630_32230 and CD630_32250), an intermediate in lysine synthesis, was predicted to sustain approximately 1% of pyruvate flux over the time course, attaining a peak of 9% pyruvate flux around 36 hours. Synthesis of 1-deoxy-D-xylulose-5-phosphate, an intermediate in thiamine synthesis, by 1-deoxy-D-xylulose-5-phosphate synthase (*dxs*, CD630_12070) was predicted to steadily decline from 0.2% pyruvate flux at the initial timepoint to nominal levels by 48 hours. Synthesis of 2-aceto-2-hydroxybutyrate from 2 pyruvate by acetolactate synthase (CD630_15660; *ilvB*) was predicted only after 38 hours, accounting for ≤0.3% of pyruvate flux at 48 hours.

To maintain nitrogen balance between Stickland and glycolytic metabolism, model predictions identified amino group transfers from Stickland oxidative leucine and valine metabolism to 2-oxoglutarate, forming glutamate. During peak metabolism, between 10-20 hours, amino groups released from Stickland amino acid fermentation in excess of biosynthetic quantities were secreted as L-alanine.

## Discussion

We illustrate novel use of HRMAS ^13^C NMR to define the complex pathways of carbohydrate and amino acid metabolism in the obligate anaerobe *C. difficile*. Obligate anaerobes live closer to thermodynamic limits of metabolism than aerobic species and utilize electron acceptors with reduced electronegative potential compared to oxygen [29, 30]. Anaerobes thus face greater constraints in coordinating simultaneous flux through multiple pathways to acquire energy and support growth while maintaining redox balance and access to usable pools of carbon, nitrogen, and electron transporters.

HRMAS NMR was performed in a media formulation with other natural abundance carbon sources to evaluate specific ^13^C carbon source metabolism in the context of co-occurring pathways of carbohydrate and amino acid fermentation. Use of heavy-tagged compounds, which can include ^15^N, ^2^H or other tags, enables detailed dissection of anaerobe metabolism within complex media conditions, including ones encountered *in vivo*.

Real-time metabolism of ^13^C-glucose identified early production of ^13^C-acetate and ^13^C-H_2_CO_3_ from the condensation of carbon dioxide with water. The HRMAS NMR data informed model predictions showing pyruvate-ferredoxin oxidoreductase (*pfo;* CD630_26820) to be the primary pathway involved in the metabolism of pyruvate to acetyl-CoA, with reduction of ferredoxin (Figure 3A). Subsequent metabolism of acetyl-CoA to acetylphosphate and acetate produces an equivalent of ATP. These reactions provide rapid sources of energy and, alone, would result in a net reductive state among pools of cellular NADH and ferredoxin. In contrast, co-occurring Stickland proline reductase metabolism, over 2.5-10 hours, provides a non-glycolytic pathway to regenerate oxidized electron carriers prior to when the reductive products of ^13^C-glucose metabolism appeared, namely ^13^C-lactate and ^13^C-ethanol. Given the presence in MMM of natural abundance carbon isotopes of the Stickland electron acceptors proline, glycine and leucine, the reductive Stickland pathways [14, 17] may provide a mechanism to regenerate oxidized NAD+ from the NADH produced in early stages of glycolytic fermentation. dFBA model predictions of pyruvate metabolism also included further transfer of ^13^C carbons into biosynthetic reactions for amino acids, cell wall components, and required vitamin cofactors for support of protein synthesis and biomass production.

By 36 hours, *C. difficile’s* metabolism shifted profoundly to reductive fermentation of glycolytic intermediates to lactate and ethanol. This shift to reductive carbohydrate fermentation followed predicted depletion of the Stickland electron acceptor amino acids proline, leucine, and glycine [31, 32]. Using HRMAS NMR we illustrate capacity to detect global, cellular shifts in integrated redox reactions using ^13^C inputs for glycolytic and Stickland pathways.

Informed by the experimental results from HRMAS ^13^C NMR, our updated *C. difficile* metabolic model predicts the formation of ^13^C-alanine from the transamination of ^13^C-pyruvate (Figure 3). While transamination of pyruvate provides no additional energetic yield, it suggests alanine’s importance as a nitrogen sink, particularly in Stickland fermenting species [26, 32]. The oxidative Stickland reactions deaminate electron-donor amino acids, with proposed transfer of the amine group to 2-oxoglutarate to form glutamate [32]. The recycling of glutamate to 2-oxoglutarate reduces NAD+, diverting pools of NAD from glycolytic and other energy-generating pathways. In contrast, transamination of pyruvate to alanine is not predicted to reduce NAD+, nor divert it from the electron carrier pool, and thus may provide a more favorable cellular sink for ammonia. Alanine also supports additional cellular functions in protein and peptidoglycan synthesis [33]. Furthermore, pools of alanine can be stored for later energy generation through oxidative Stickland metabolism, or via mixed acid fermentation reactions with alanine’s deamination to pyruvate [16, 31].

Informed by experimental findings from HRMAS NMR the metabolic model predicts ferredoxin to be the primary electron acceptor in both carbohydrate and amino acid fermentative pathways, and NADH to be the primary electron donor. Ferredoxin’s reduction occurs primarily via the oxidative decarboxylation of 2-oxocarboxylic acids by *pfo* and by the oxidative Stickland reactions. In contrast, model predictions indicate NADH oxidation to occur via the Wood-Ljungdahl pathway throughout the time course, with additional significant NADH oxidation from flux through the proline reductase pathways over early metabolism during the first 12 hours and then from mixed acid fermentation and glycine reductase flux at ≥30 hours. Model predictions further indicate that the Rnf complex provides a primary system of electron transfer from reduced ferredoxin to NAD^+^ while also playing a critical role in cellular ATP generation [16, 34]. By maintaining a net reductive state of ferredoxin and net oxidative state of NAD+ through coordinate fermentation of carbohydrates and amino acids, *C. difficile* augments its capacity to generate proton gradients through the Rnf complex, an energy source which can be harnessed for ATP production, cellular motility, and other functions. These findings identify alanine, and the pathways modulating its production, as new targets to potentially combat *C. difficile* infections [35].

We present the first high-resolution studies of real-time metabolism in living obligately anaerobic cells and show new integration points to maintain nitrogen and redox balance. Integrated use of HRMAS ^13^C NMR with metabolic modeling offers a greatly improved approach to define cellular metabolism and integration points within obligate anaerobes to support clinical, scientific and industrial applications.

## Materials and Methods

### Strains

A PaLoc-deleted strain of ATCC 43255 was generated that lacked the *tcdB, tcdE* and *tcdA* genes to reduce biohazard risks for NMR analyses. The deletion mutant was created using a recently developed toxin-mediated allele exchange method [36]. Briefly, two regions of approximately 800 bp of DNA flanking the region to be deleted were amplified by PCR from *C. difficile* ATCC43255 using the primers in Supplemental Table 3. Purified PCR products were cloned into the PmeI site of the pMSR0 vector using NEBuilder HiFi DNA Assembly. The resulting plasmid was transformed into *E. coli* strain NEB10β (New England Biolabs) and the insert verified by sequencing. *E. coli* strains were cultured aerobically at 37 °C in LB broth or LB agar supplemented with chloramphenicol (15 μg/ml). The plasmid was then transformed into *E. coli* HB101 (RP4) and conjugated into *C. difficile* ATCC43255 that had been heat-shocked at 50°C for 15 min beforehand [37]. Transconjugants were selected on BHI agar plates supplemented with cycloserine (250 μg/ml), cefoxitin (25 μg/ml), and thiamphenicol (15 μg/ml). Allelic exchange was performed according to Peltier et al. [36]. This strain was shown to be non-toxigenic to human fibroblasts and gnotobiotic mice (data not shown).

### Strain culture conditions

The PaLoc-strain of ATCC 43255 was cultured for 12 hours in supplemented brain-heart infusion media (BHIS; Remel, Lenexa, KS). Cells were spun and washed three times in pre-reduced PBS, prepared in molecular-clean water, and diluted to introduce 100,000 cells into HRMAS NMR rotor inserts for analyses. Preparations were serially diluted and plated to Brucella agar (Remel, Lenexa, KS) to quantitate vegetative cells and spores used in input preparations. Spore counts after 12 hours of culture in BHIS were <0.1% of vegetative cells.

*C. difficile* Modified Minimal Medium (MMM) was prepared as described [38] with U-^13^C glucose or U-^13^C proline as indicated in Table 2. Conditions evaluating the effects of exogenous selenium added 0.1uM sodium selenite to the media (Sigma Chemical, St. Louis, MO).

**Table 2.** Media conditions for HRMAS ^13^C NMR.

| Medium Component | Concentration |  |  |
| --- | --- | --- | --- |
| | U-$^{13}\text{C}$ Glucose | U-$^{13}\text{C}$ Proline + Se | U-$^{13}\text{C}$ Proline – Se |
| U-$^{13}\text{C}$ Glucose* (mM) | 27.5mM | 0 | 0 |
| U-$^{13}\text{C}$ Proline** (mM) | 0 | 30mM | 30mM |
| Sodium Selenite ($\mu\text{M}$) | 0.1uM | 0.1uM | trace*** |

\* $^{13}\text{C}$ proline condition formulations contain 11 mM unlabeled glucose, as specified in the MMM formulation. $^{13}\text{C}$ glucose condition formulation does not contain unlabeled glucose.
\*\* $^{13}\text{C}$ glucose and $^{13}\text{C}$ proline condition formulations contain 5.22 mM unlabeled proline from the MMM formulation.
\*\*\*Selenium-limiting condition contained trace amounts of selenium from impurities in other media components.

NMR rotors were loaded in an anaerobic chamber with 100,000 CFU added to the defined MMM conditions. The insert was sealed and removed from the chamber for NMR analyses.

After analyses, rotor contents were checked for pH, serially diluted and plated to Brucella agar and incubated anaerobically at 37°C to evaluate vegetative and spore CFU and confirm absence of contaminating species. Cellular growth in the rotor was approximately 30% over the input biomass. Given buffering components in the media, the pH did not drop below 7.0 in any conditions after 48 hours of analyses.

### HRMAS NMR

HRMAS NMR measurements were performed on a Bruker Avance (Billerica, MA) 600 MHz spectrometer. The Kel-F insert loaded in the anaerobic chamber was placed in a 4 mm zirconia rotor with 2 uL of D2O added for field locking before the rotor was sealed and introduced into the triple-tuned HRMAS probe. One and two dimensional ^1^H and ^13^C NMR were conducted at 37 °C with a spin-rate of 3600 ± 2 Hz. One-dimensional time series spectra were measured alternately and continuously for ^1^H NOESY with water suppression (~13 mins) and for proton decoupled ^13^C (~43 mins) throughout the length of the experimental time. Two-dimensional ^1^H COSY (~3 hrs and 49 mins), proton decoupled ^13^C COSY (~3 hrs and 30 mins), and ^1^H-^13^C HSQC (~3 hrs and 38 mins) spectra were inserted in the between the 1D time series. MRS spectra were processed using the TopSpin 3.6.2 (Billerica, MA).

### ^13^C and proton spectra analyses

^13^C spectral signal tables from each time point were concatenated with a list of reference ppm into a combined matrix (Supplemental File 1). Further processing was performed in MATLAB R2019b (MathWorks, Natick, MA). Spectra were normalized by noise root-mean-square error (RMSE) of the 140–160 ppm region. The normalized spectrum stack was visualized as a surface plot with face lightness mapped to the log base-2 of signal. Local maxima within 0.5 ppm of each reference peak with height ≥4 times the noise RMSE were classified as detectable signals and concatenated into signal ridges. Signal ridges were color-labeled by metabolite and superimposed over the stack as a scatter plot with stems to the *xy* plane and a smoothing spline curve fit. Surface regions within 0.5 ppm of each reference peak were colorized.

Logistic curve fits for each compound in the time series used the following equation:

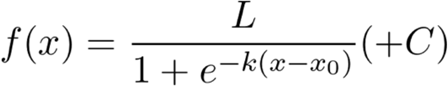

*L*: Maximum value of the logistic curve
*k:* Growth rate of the curve
*x*_0_: *x* value of the mid-point of the sigmoidal curve
*C*: Signal value of the input U-^13^C carbon source at 48 hours

Average peak heights for each compound were plotted and the logistic curve fit performed in MATLAB. The custom python and MATLAB scripts are available via GitHub (https://github.com/Massachusetts-Host-Microbiome-Center/nmr-cdiff).

### Metabolic modeling

A previously published genome-scale metabolic model of *C. difficile* strain 630, icdf834 [28] was converted into COBRA json format using the COBRA toolbox in MATLAB [39]. The model was modified using the COBRApy toolbox [40] and custom python scripts. Exchange reaction bounds were set to 3% of the millimolar concentration of media components used experimentally [38]. The following changes were made based on experimental data and to support biologically relevant processes energetically and thermodynamically.

1. An exchange reaction was added for iron(II) and transport and secretion reactions were added for propionate, phenylacetate, indole-3-acetate, butyrate, and n-butanol.
2. Reactions of the oxidative Stickland pathways were added, employing ferredoxin as the oxidizing agent (Supplemental Table 1, rows 1-3), according to previous descriptions of electron bifurcation in Stickland metabolism [16, 41].
3. Additionally, a proton motive force (PMF)-generating Rnf complex reaction and an ATP synthase reaction were added (Supplemental Table 1,, rows 4-5) to complete the Stickland energy salvage mechanism.
4. Proposed pathways for butyrate and a putative propionate fermentation were added (Supplemental Table 1, rows 6-12) based on experimental evidence of secretion in *C. difficile* [32, 42].
5. Reversibility of some existing reactions were altered to reflect practical thermodynamic bounds and prevent cycles from occurring in the flux balance analysis (FBA) solution (Supplemental Table 1, rows 13-24).
6. Reactions for free H+ transport and thioredoxin oxidation were removed because they were thermodynamically infeasible; succinyl-CoA synthase and putrescine transaminase reactions were removed due to a lack of genomic evidence (Supplemental Table 1, rows 25-28).

### Dynamic Flux Balance Analysis (dFBA)

The primary mechanisms of ^13^C relaxation in small molecules are dipole-dipole interactions and spin-rotation [43]. Thus, the signal intensity of a ^13^C atom is highly dependent on its molecular context, especially regarding the atom’s attachments to ^1^H atoms and its ability to freely rotate. With the assumption that ^13^C atoms in equivalent molecular contexts are subject to similar relaxation times, the relative abundances of ^13^C metabolites as analyzed by HRMAS NMR were estimated by comparing the integrated signal of carbon atoms in homologous functional groups. The integrated signals of glucose C-6 at the initial timepoint and ethanol C-1 at the final timepoint were selected for comparison due to their shared –CH_2_OH context. The final concentration of ethanol was estimated by multiplying the initial concentration of glucose by the ratio of integrated ethanol C-1 signal to integrated glucose C-6 signal. Final concentrations of acetate, alanine, and lactate were estimated by multiplying the ethanol concentration by the ratio of acetate C-2, alanine C-3, and lactate C-3 integrated signal, respectively, to ethanol C-2 integrated signal, due to the common –CH3 molecular context. Additionally, the final concentration of 5-aminovalerate was estimated by multiplying the initial concentration of proline by the ratio of proline C-4 integrated signal at the initial timepoint to 5-aminovalerate C-3 integrated signal at the final timepoint.

dFBA was implemented by computing steady-state flux balance analysis (FBA) solutions with time-dependent exchange fluxes using the COBRApy toolbox [30] and custom python scripts. Metabolite concentration trajectories were estimated by computing a logistic fit on the integrated ^13^C signal and scaling the logistic function by the estimated relative abundance. Exchange (for glucose and proline) and secretion (for end metabolites) upper and lower bounds were set to the derivative of the concentration logistic function, evaluated per time point. Steady-state FBA solutions were calculated along a simulated 48-hour timescale with a resolution of 1 solution per hour.

## Acknowledgments

We thank Michael Judge, Arthur Edison, Mario Arietta-Ortiz, Nitin Baliga and Linc Sonenshein for critical reading of the manuscript and helpful comments.

