## Supplemental Figures and Tables for "Real-time HRMAS ^13^C NMR of obligately anaerobic cells identifies new metabolic targets in the pathogen *Clostridioides difficile*"

**Supplemental Figure 1—$^1$H-$^{13}$C 2D NMR spectrum of $^{13}$C glucose metabolites.** Metabolites were identified from $^1$H-$^{13}$C 2D HSQC NMR spectra using reference spectra from HMDB.

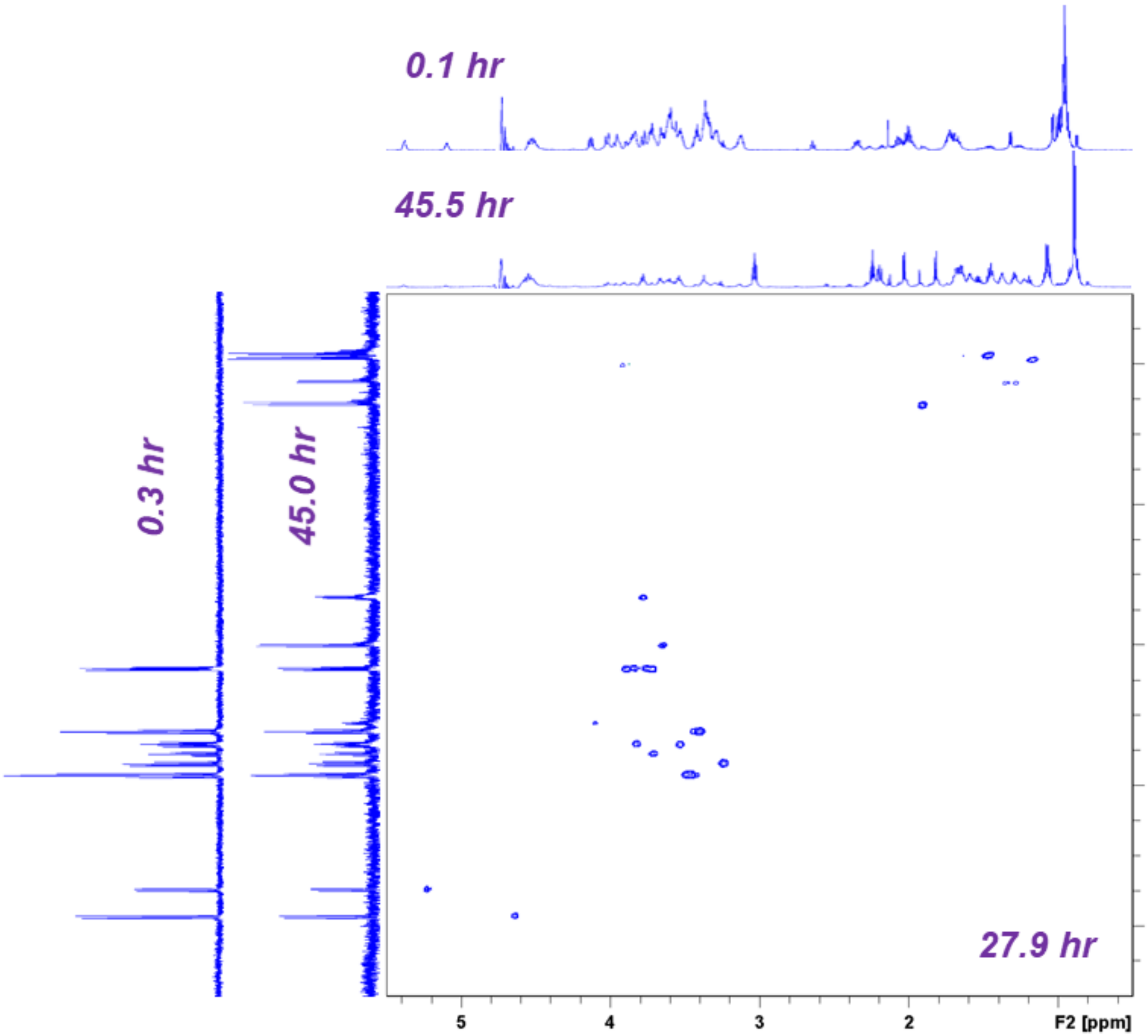

**Supplemental Figure 2—$^1$H-$^{13}$C 2D NMR spectrum of $^{13}$C proline metabolites.** Metabolites were identified from $^1$H-$^{13}$C 2D HSQC NMR spectra using reference spectra from HMDB.

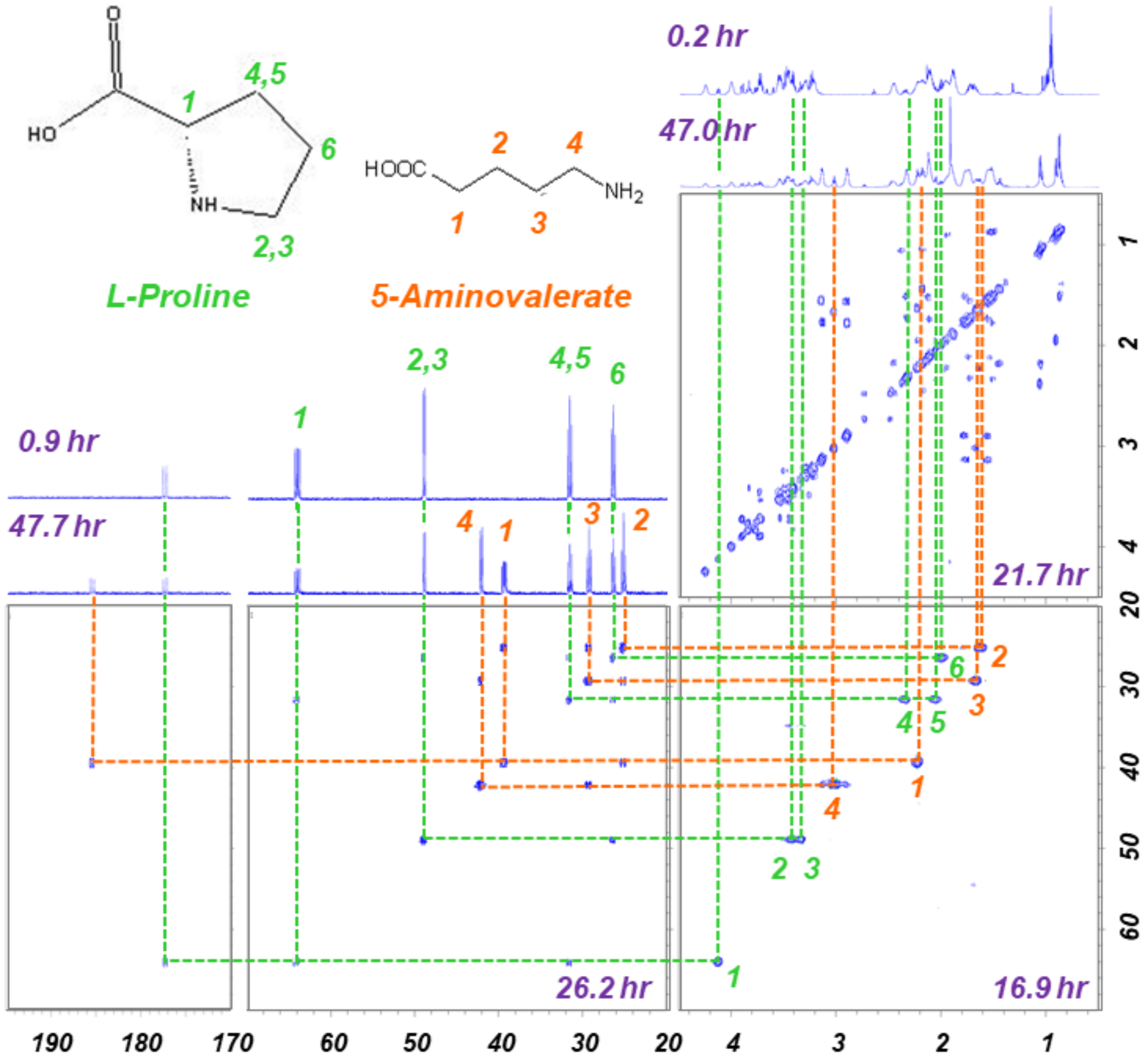

### Supplemental Table 1—Major fermentative pathway genes of *C. difficile.*

| Fig 3 Label | System Name | Gene Association | |
|---|---|---|---|
| | | **Strain CD630** | **Strain ATCC43255** |
| 1 | Glucose PTS transport | (CD630_03880) or (CD630_31270) or (CD630_31250) or (CD630_30970) or (CD630_30300 and CD630_30270) | (UAB_RS02000000220100) or (UAB_RS02000000220895) or (UAB_RS0216690) or (UAB_RS0216545) or (UAB_RS0216180 and UAB_RS0216165) |
| | Glucose 6-phosphotransferase | CD630_24590 or CD630_22360 | UAB_RS0213555 or UAB_RS0212390 |
| | G6P ketol-isomerase | CD630_32850 | UAB_RS0217935 |
| | Phosphofructokinase (PFK) | CD630_31100 or CD630_33950 | UAB_RS0216610 or UAB_RS0218270 |
| | 1,6-FBP aldolase | CD630_04030 or CD630_31350 | UAB_RS0203140 or UAB_RS0216740 |
| | Triose-phosphate isomerase | CD630_31720 | UAB_RS0217175 |
| | G3P dehydrogenase | CD630_31740 or CD630_17670 | UAB_RS0217185 or UAB_RS0210085 |
| | Phosphoglycerate kinase | CD630_31730 | UAB_RS0217180 |
| | G3P:NADP+ oxidoreductase | CD630_05800 | UAB_RS0203980 |
| | Phosphoglycerate mutase | CD630_31710 or CD630_18140 | UAB_RS0217170 or UAB_RS0210340 |
| | Phosphopyryvate hydratase | CD630_31700 | UAB_RS0217165 |
| | Pyruvate kinase | CD630_33940 | UAB_RS0218265 |
| 2 | Proline racemase | CD630_32370 | UAB_RS0217570 |
| | Proline Reductase *prd* | CD630_32400 and CD630_32380 and CD630_32370 and CD630_32410 and CD630_32430 and CD630_32440 | UAB_RS0217585 and UAB_RS0217575 and UAB_RS0217570 and UAB_RS2000000221565 and UAB_RS0217600 and UAB_RS0217605 |
| | Glycine Reductase *grd* | CD630_23540 and CD630_23520 and CD630_23510 and CD630_23490 and CD630_23480 and CD630_23570 | UAB_RS0213000 and UAB_RS0212990 and UAB_RS0212985 and UAB_RS0212975 and UAB_RS0212970 and UAB_RS0213015 |
| | 4-methyl-2-oxopentanoate reductase *ldhA* | CD630_03940 | UAB_RS0203090 |
| | Isocaprenoyl-CoA:2-hydroxyisocaproate CoA-transferase *hadA* | CD630_03950 | UAB_RS0203095 |
| | 2-hydroxyglutaryl-CoA dehydratase *hadBC* | CD630_03970 and CD630_03980 | UAB_RS0203105 and UAB_RS0203110 |
| 3 | 2-oxoacid dehydrogenase (ferredoxin) | (CD630_01150 and CD630_01160 and CD630_01170 and CD630_01180) or | (UAB_RS0201425 and UAB_RS0201430 and |

| | | | |
|---|---|---|---|
| | | (CD630_21970 and CD630_21980 and CD630_21990 and CD630_21991) or (CD630_24270 and CD630_24280 and CD630_24290 and CD630_24291) | UAB_RS0201435 and UAB_RS0201440) or (UAB_RS0212180 and UAB_RS0212185 and UAB_RS0212190 and UAB_RS0212195 or (UAB_RS02000000220775 and UAB_RS0213385 and UAB_RS0213390 and UAB_RS0213395) |
| | Acyl-CoA thioesterase | CD630_01120 or CD630_19660 | UAB_RS0201410 or UAB_RS0210940 |
| | Acyl kinase | CD630_11750 or CD630_23790 or CD630_24260 | UAB_RS0206515 or UAB_RS0213135 or UAB_RS0213375 |
| 4 | Amino acid:2-oxoglutarate aminotransferase | CD630_01070 or CD630_13390 or CD630_28280 or CD630_15490 or CD630_22000 or CD630_21580 | UAB_RS0201370 or UAB_RS0207590 or UAB_RS0215480 or UAB_RS0208695 or UAB_RS0212200 or UAB_RS0211975 |
| 5 | Glutamate dehydrogenase | CD630_15370 | UAB_RS0208625 |
| 6 | Alanine transaminase | CD630_13390 | UAB_RS0207590 |
| 7 | Rnf complex | CD630_11370 and CD630_11380 and CD630_11390 and CD630_11400 and CD630_11410 and CD630_11420 | UAB_RS0206325 and UAB_RS0206330 and UAB_RS0206335 and UAB_RS0206340 and UAB_RS0206345 and UAB_RS0206350 |
| 8 | F-type ATP synthase | CD630_34700 and CD630_34740 and CD630_34720 and CD630_34710 and CD630_34690 and CD630_34680 and CD630_34730 and CD630_29600 and CD630_34760 | UAB_RS0218655 and UAB_RS0218675 and UAB_RS0218665 and UAB_RS0218660 and UAB_RS0218650 and UAB_RS0218645 and UAB_RS0218670 and UAB_RS0215835 and UAB_RS0218685 |
| 9 | Pyruvate:ferredoxin oxidoreductase | CD630_01180 and CD630_21990 and CD630_21980 | UAB_RS0201440 and UAB_RS0212195 and UAB_RS0212185 |
| 10 | Pyruvate formate lyase (*pfl*) | CD630_07590 or CD630_11200 or CD630_32820 | UAB_RS0204945 or UAB_RS0206220 or UAB_RS0217920 |
| 11 | Formate dehydrogenase (*fdhF*) | CD630_33170 | UAB_RS0218100 |
| 12 | Acyl-CoA thioesterase | CD630_19660 | UAB_RS0210940 |
| | Acetate kinase | CD630_11750 | UAB_RS0206515 |
| 13 | Alcohol dehydrogenase | CD630_03340 or CD630_30060 or CD630_29660 or CD630_19170 | UAB_RS0202955 or UAB_RS0216065 or UAB_RS0215865 or UAB_RS02000000220555 |
| Not in Fig3 | Serine-pyruvate transaminase | CD630_09940 | UAB_RS0205710 |
| Not in Fig3 | Acetolactate synthase | CD630_15660 | UAB_RS0208775 |

| | | | |
|---|---|---|---|
| Not in Fig3 | 4-hydroxy-tetrahydrodipicolinate synthase | CD630_32230 and CD630_32250 | UAB_RS0217490 and UAB_RS0217500 |
| Not in Fig3 | 1-deoxy-D-xylulose-5-phosphate synthase | CD630_12070 | UAB_RS0206680 |

### Supplemental Table 2—Pathway modifications to the icdf834 model

| Row # | Reaction ID | Reaction name | Reaction | Modification |
|---|---|---|---|---|
| 1 | RXN-19534 | 3-methyl-2-oxobutanoate dehydrogenase (ferredoxin) | 4m2op_c + coa_c + 2 feroxoxi_c <=> co2_c + 2 feroxred_c + h_c + ivalcoa_c | Added |
| 2 | ICCoA-DHG-EB | 2-isocaprenoyl-CoA dehydrogenase (electron-bifurcating) | 2 feroxoxi_c + isocaprecoa_c + 2 nadh_c <=> 2 feroxred_c + isocapcoa_c + 2 nad_c | Added |
| 3 | CPLX-8556 | 3-methyl-2-oxobutanoate dehydrogenase (ferredoxin) | 2oiv_c + coa_c + 2 feroxoxi_c <=> co2_c + 2 feroxred_c + h_c + isobutcoa_c | Added |
| 4 | RNF-Complex | RNF Complex (PMF-generating) | 2 feroxred_c + h_c + nad_c --> 2 feroxoxi_c + nadh_c + 2 pmf_c | Added |
| 5 | ATPsynth4_1 | ATP synthase 4:1 | adp_c + pi_c + 4 pmf_c --> atp_c | Added |
| 6 | CrotCoA_DHG_EB | Crotonyl-CoA dehydrogenase (electron-bifurcating) | ctncoa_c + 2 feroxoxi_c + 2 nadh_c <=> butcoa_c + 2 feroxred_c + 2 nad_c | Added |
| 7 | PBT | Phosphate butyryltransferase | butcoa_c + pi_c --> butp_c + coa_c | Added |
| 8 | BUK | Butyrate kinase | adp_c + butp_c --> atp_c + but_c | Added |
| 9 | ADH_ButCoA | Alcohol dehydrogenase Butyryl-Coa:Butanal | butcoa_c + h_c + nadh_c <=> btal_c + coa_c + nad_c | Added |
| 10 | ADH_Btal | Alcohol dehydrogenase Butanal:Butanol | btal_c + h_c + nadh_c <=> buta_c + nad_c | Added |
| 11 | PPAKr | Propionate kinase | adp_c + ppap_c <=> atp_c + ppa_c | Added |
| 12 | PTA2 | Phosphate acetyltransferase | pi_c + ppcoa_c <=> coa_c + ppap_c | Added |
| 13 | ID_28 | L-Leucine:2-oxoglutarate aminotransferase | 4m2op_c + gluL_c <=> 2oglut_c + leuL_c | Made reversible |
| 14 | ID_523 | acetolactate decarboxylase | 2alac_c --> actnR_c + co2_c | Made irreversible |
| 15 | ID_191 | N-carbamoul-L-aspartate hydrolase | aspL_c + co2_c + nh3_c <-- casp_c + h2o_c | Made irreversible |
| 16 | ID_449 | 2-acetolactate pyruvate-lyase (carboxylating) | 2.0 pyr_c --> 2alac_c + co2_c | Made irreversible |
| 17 | ID_516 | 3-phosphonopyruvate carboxy-lyase | 3ppyr_c --> co2_c + ppacal_c | Made irreversible |
| 18 | ID_303 | Phosphonoacetate phosphonohydrolase | h2o_c + ppac_c --> ac_c + pi_c | Made irreversible |
| 19 | ID_565 | oxaloacetate carboxy-lyase | oxac_c --> co2_c + pyr_c | Made irreversible |
| 20 | ID_16 | UTP phosphohydrolase | h2o_c + utp_c --> pi_c + udp_c | Made irreversible |
| 21 | Trans_h3 | sodium ion:proton antiporter | h_c + na_e --> h_e + na_c | Made irreversible |
| 22 | ID_622 | D-glyceraldehyde 3-phosphate:NADP+ oxidoreductase | glyald3p_c + h2o_c + nadp_c --> 3pg_c + h_c + nadph_c | Made irreversible |

| | | | | |
|---|---|---|---|---|
| **23** | ID_518 | 2-methylbutanoyl CoA transferase | 2mbcoa_c + pi_c --> 2mbutp_c + coa_c | Made irreversible |
| **24** | ID_539 | propanoyl-CoA:formate C-propanoyltransferase | 2obut_c + coa_c --> for_c + ppcoa_c | Made irreversible |
| **25** | ID_310 | succinyl-CoA synthase | atp_c + coa_c + succ_c --> adp_c + pi_c + succoa_c | Removed |
| **26** | Trans_h | H+ Transport | h_e --> h_c | Removed |
| **27** | ID_612 | thioredoxin oxydation | thioredoxred_c --> 2.0 h_c + thioredoxoxi_c | Removed |
| **28** | ID_295 | putrescine aminotransferase | ptrc_c + pyr_c --> 4abald_c + alaL_c | Removed |

**Supplemental Table 3—Primers used for construction of the PaLoc- strain**

| Primer | Sequence (5’ to 3’)* | Use |
|---|---|---|
| BD013 | ttttttgttaccctaagtttGGATGATTTTATGCAAAAGTC | 5’ left arm for *tcdBEA* deletion |
| BD014 | tattttagccCATAAAATTTTCTCCTTTACTATAATATTTTTAC | 3’ left arm for *tcdBEA* deletion |
| BD015 | aaattttatgGGCTAAAATATATGTTTGATAAAAAATTATTC | 5’ right arm for *tcdBEA* deletion |
| BD016 | agattatcaaaaaggagtttCCAGCTTGTTCTGAAGAC | 3’ right arm for *tcdBEA* deletion |
| BD017 | GGAGGATATATAAAAGAGTTTTAGC | 5’ screening of *tcdBEA* deletion |
| BD018 | GGGTATTGCTCTACTGGC | 3’ screening of *tcdBEA* deletion |

*Lowercase bases indicate overlapping sequences
